# Nanos downregulates maternal mRNAs in germline during *Drosophila* early embryogenesis

**DOI:** 10.1101/2025.08.18.670776

**Authors:** Yasuhiro Kozono, Makoto Hayashi, Miho Asaoka, Satoru Kobayashi

## Abstract

**Background:** Many maternal mRNAs in *Drosophila* primordial germ cells (PGCs) are degraded in concert with the synthesis of new transcripts from the zygotic genome during gastrulation and germ band elongation. However, few studies have focused on maternal mRNA destabilization in PGCs at the blastoderm stage that is prior to zygotic genome activation (ZGA). Thus, the stability of maternal mRNAs at this stage and regulation of their degradation remain poorly understood. To address this gap, we examined the role of Nanos, an RNA-binding protein known to promote mRNA degradation, in blastoderm-stage PGCs.

**Results:** By combining flow cytometry and RNA-seq analysis of PGCs, we identified the transcripts of 898 genes that were increased in *nanos*^−^ PGCs. Among them, maternal mRNAs encoded by 298 genes were downregulated by Nanos in PGCs.

**Conclusions:** Our results show that Nanos downregulates maternal mRNA expression in PGCs before ZGA. As Nanos in *C. elegans* PGCs has also been reported to promote maternal-to-zygotic transition (MZT) via maternal mRNA downregulation during a transcriptionally silent state, our findings highlight the importance of investigating the function of Nanos for understanding the MZT in PGCs across various animal species.

## 1 INTRODUCTION

During the maternal-to-zygotic transition (MZT), maternal mRNAs are degraded in concert with the synthesis of new transcripts from the zygotic genome. Several studies have shown that delays in their degradation impedes proper embryonic development^1,2^, indicating that maternal mRNA decay must be tightly regulated.

In many animals, primordial germ cells (PGCs) are specified by inheriting factors localized in a specialized ooplasm, or germ plasm, which are required for their development. In *Drosophila*, PGCs are formed by incorporating the germ plasm, which harbors maternal mRNAs, and resides at the posterior pole of the blastoderm [1.5–3 h after egg laying (AEL)]. Massive maternal mRNA degradation and zygotic genome activation (ZGA) in PGCs during gastrulation and germ band elongation (3–5 h AEL) were first observed through transcriptomic and genetic analyses^1,3,4^. However, as few studies have focused on mRNA destabilization in blastoderm stage PGCs, maternal mRNA regulation in germline before ZGA remains poorly understood.

Nanos is an evolutionarily conserved RNA-binding protein that promotes degradation and/or translational repression of mRNAs^5–7^. In *Drosophila*, Nanos forms a complex with the RNA-binding protein Pumilio (Pum), and binds to the Nanos–Pum motif, defined as WWWUGUA (W = A/U)^8^. During embryogenesis, Nanos is produced at the posterior pole immediately after egg deposition, transiently distributed in abdominal region at cleavage stage, and restricted to PGCs at blastoderm stage^9^. Nanos represses translation of *hunchback* (*hb*) mRNA in somatic regions, and of *Cyclin B* (*CycB*) and *importin α2* (*impα2*) mRNAs in blastoderm-stage PGCs^10–14^. The repression of *CycB* and *impα2* are necessary for PGC development^10,12,15,16^. In addition, Nanos would broadly regulate maternal mRNAs. RNA-seq was performed on embryos harboring a fusion protein of Nanos and the nonsense-mediated decay (NMD) factor Upf1^17^. Given that tethering Upf1 to mRNAs induces their degradation. Using this system, they identified approximately 2,600 maternal mRNAs that were downregulated by Upf1-Nanos (Upf1-Nanos targets)^17^, indicating that these mRNAs are deposited maternally and potentially regulated by Nanos.

Here, to identify maternal mRNAs that are actually downregulated by Nanos in PGCs, we purified PGCs from normal and *nanos* loss-of-function (*nanos*^*-*^) embryos at 2–3 h AEL, when the massive degradation of maternal mRNAs is not thought to have occurred, and performed RNA-seq analysis of these PGCs.

## 2 RESULTS AND DISCUSSION

### 2.1 Maternal mRNAs encoded by 298 genes were downregulated by Nanos in PGCs

We performed RNA-seq analysis of PGCs from normal and *nanos*^−^ embryos (Figure 1A). *nanos* mRNA was significantly reduced in *nanos*^−^ PGCs compared to that in normal PGCs, demonstrating the reliability of our RNA-seq data. The mRNAs encoded by 898 genes were significantly increased in *nanos*^−^ PGCs (Table S1). These mRNAs are thus downregulated in a Nanos-dependent manner. Among them, we found that maternal mRNAs encoded by 298 genes (Group I) were in the list of Upf1-Nanos targets (Table S2), while those encoded by 600 genes (Group II) were not.

**Figure 1.**
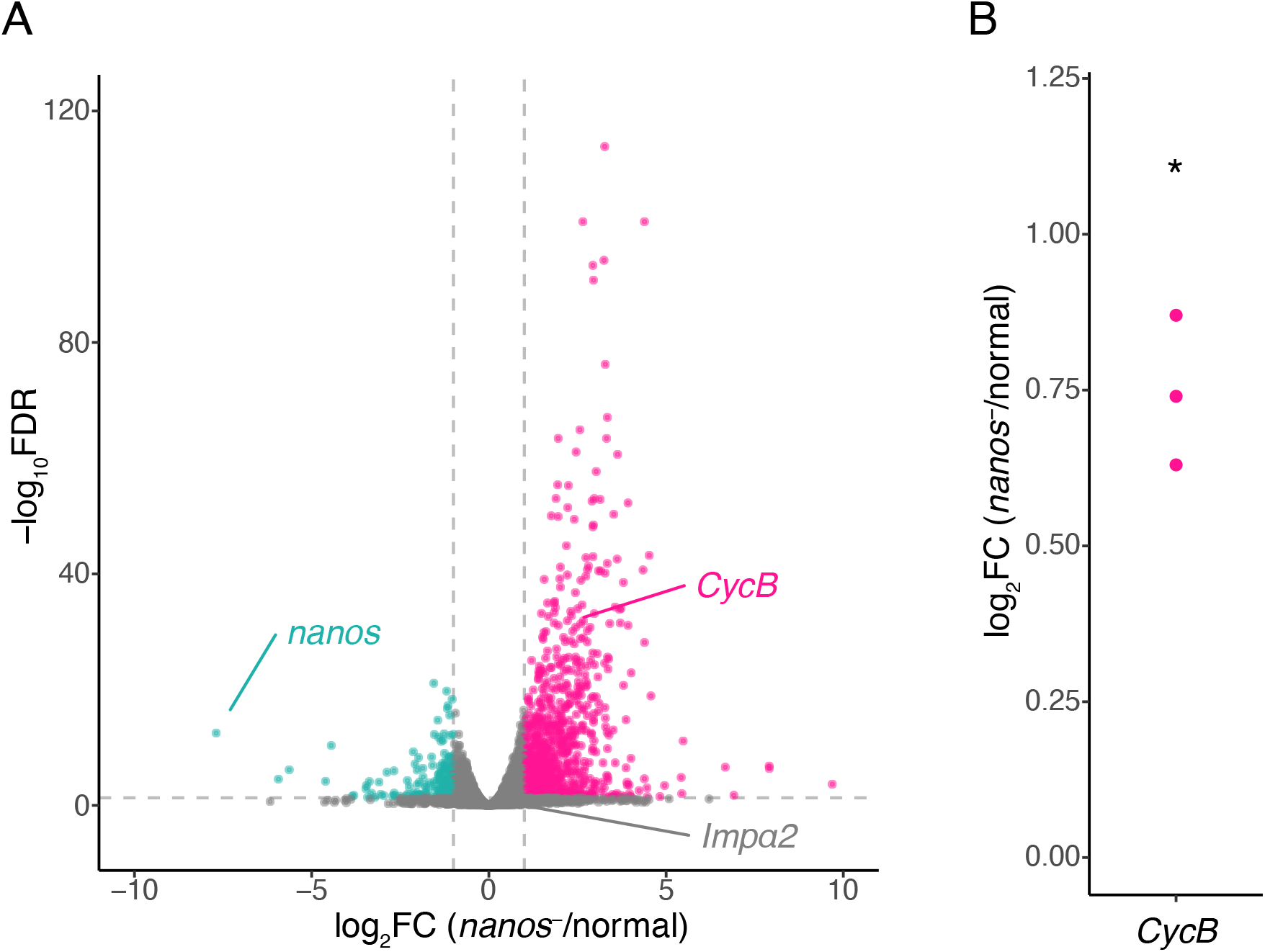
Transcriptome and RT-qPCR analyses for *nanos-*PGCs. (A) Volcano plot for normal vs *nanos-* PGCs. The thresholds for statistical significance were defined as FDR < 0.05 and log2FC ≥ 1 or ≤ −1. Magenta dots indicate the significantly increased mRNAs in *nanos-* PGC, and cyan dots indicate significantly decreased mRNAs. *CycB* mRNA was significantly increased in *nanos*-PGCs, while *impα2* mRNA was not. (B) Log2 fold changes (log2FC) in *nanos-*/normal of *CycB* mRNA are presented. Significance was calculated by paired *t*-test. *P < 0.05.

### 2.2 The role of “Group I” mRNAs on PGC development

To gain insights into Nanos function, we performed Gene Ontology (GO) analysis for “Group I” mRNAs (Table S3). Maternal mRNAs were enriched in terms related to the regulation of “GTPase activity”, suggesting that Nanos might play a role in controlling signal transduction. This GO category included somatic genes such as *RhoGAP15B* and *Asap*, which are involved in imaginal disc-derived leg morphogenesis and eye morphogenesis, respectively^18,19^. These results suggest that Nanos play a role in downregulation of somatic maternal mRNAs. A previous study has shown that *nanos*^*-*^ PGCs can differentiate into somatic cells^20^; this may be accounted for Nanos-mediated down regulation of somatic maternal mRNAs.

### 2.3 The enrichment of Nanos-Pum motif in both “Group I and II” mRNAs

To know whether the “Group I” mRNAs are enriched for the binding motif of Nanos-Pum, we focused on Nanos-Pum motif, defined as WWWUGUA (W = A/U). This is motivated by several previous observations: (1) Nanos binds to the Nanos-Pum motif cooperatively with Pum^8^, based on SEQRS (*in vitro* selection, high-throughput sequencing of RNA, and sequence specificity landscapes) analyses findings; (2) partial deletion of the Nanos-Pum motif (AUUGUA) from *hb* 3′-UTR abolishes Nanos-dependent mRNA degradation^6^, and (3) Upf1-Nanos targets are enriched for this motif^17^. We first identified Nanos-downregulated mRNA isoforms in PGCs because several mRNA isoforms are transcribed from a single gene. When we reanalyzed our data, 491 mRNA isoforms were found to be significantly upregulated in *nanos*^−^ PGCs (Table S4). We next analyzed the proportion of mRNA isoforms that contain at least one Nanos-Pum motif and its density (the number of motifs per kilobase of 3′ UTR)^17^. As expected, we found that the “Group I” mRNAs showed significantly higher motif rates and densities compared to mRNAs that were not significantly altered in the presence and absence of Nanos (Figure 2). Unexpectedly, this enrichment was also observed in “Group II” (Figure 2), suggesting that these mRNAs may contain maternal transcripts regulated by Nanos in PGCs. This notion is supported by the fact that “Group II” contains *CycB* mRNA targeted by Nanos. *CycB* mRNA is known to be bound and translationally repressed by Nanos^8,10,11^. Furthermore, *CycB* mRNA is downregulated by Nanos in PGCs in our RNA-seq and RT-qPCR analyses (Figure 1).

**Figure 2.**
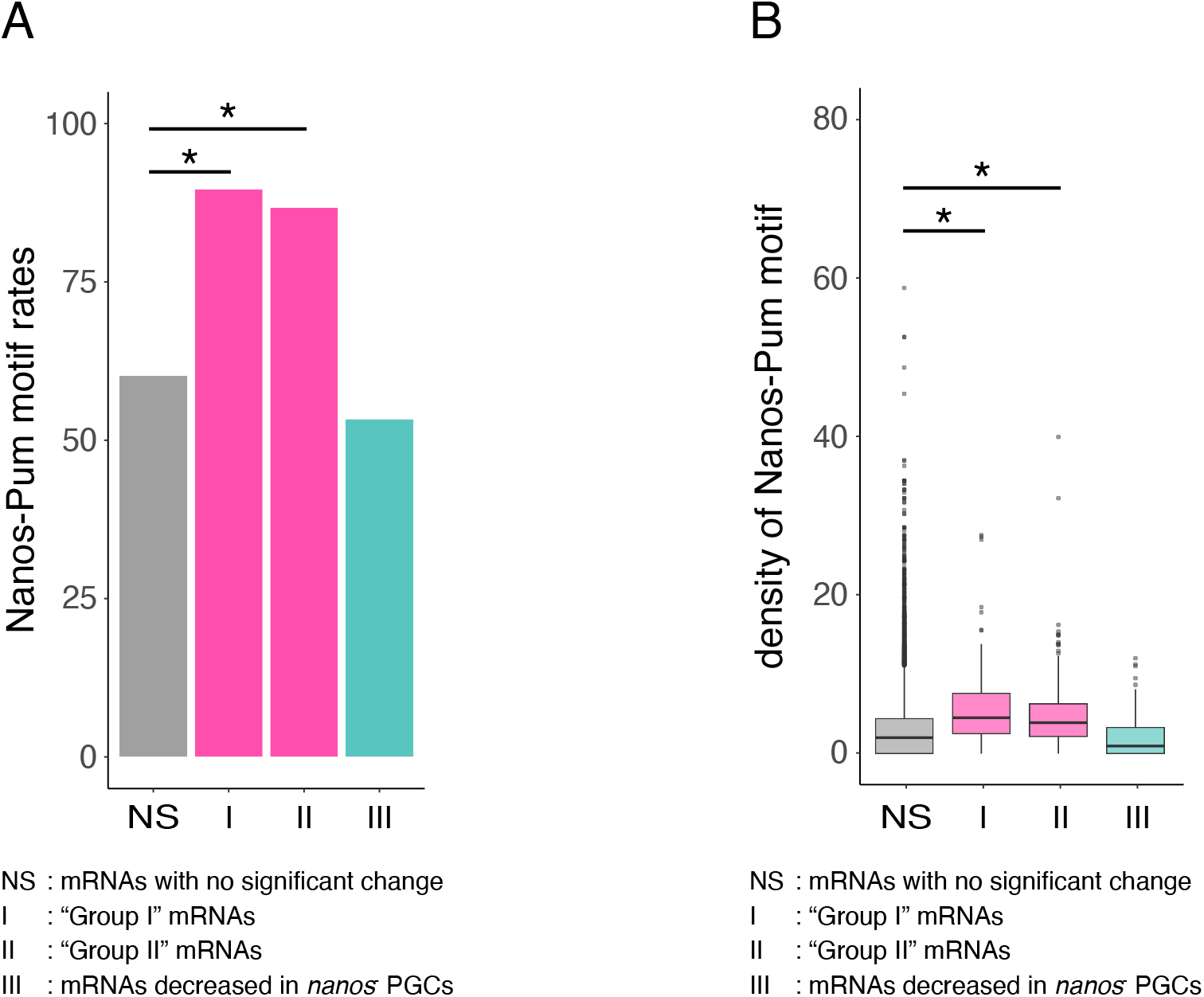
The analyses Nanos-Pum motif. (A) the proportion of mRNAs that contain at least one Nanos-Pum motif was shown. NS: mRNAs with no significant change; I: “Group I” mRNAs; II: “Group II” mRNAs; III: mRNAs decreased in *nanos*^−^ PGCs. Significance was calculated using Fisher’s exact test with Bonferroni correction. *P < 0.05. (B) The density of this motif (number per kilobase of 3′ UTR of this motif) was shown. Labels of each group were the same as in (A). Significance was assessed using the Steel–Dwass test. *P < 0.05.

### 2.4 Germline-specific function of Nanos in maternal mRNA destabilization

The *hb* mRNA was included in “Group I” mRNAs (Table S2). Given that *hb* mRNA is also downregulated by Nanos in the somatic region^17^, it is likely that *hb* mRNA is regulated through a common mechanism in both soma and PGCs. In contrast, stability of *CycB* mRNA may be regulated through distinct mechanisms in soma and PGCs. Our results indicate that Nanos downregulates *CycB* mRNA in PGCs. However, previous RNA-seq analyses comparing normal and *nanos*^−^ embryos did not detect significant differences in *CycB* mRNA levels^17^. Furthermore, it is suggested that *CycB* mRNA in somatic region is downregulated in a Smaug-dependent manner^21^. Smaug is a major factor for the clearance of maternal mRNAs in the soma^1,22^, and consistent with this, *smaug* mutants block the downregulation of *CycB* mRNA level^21^.

This is the first study to report that Nanos downregulates a wide range of maternal mRNAs during early germline development. Immediately following the analyzed stage, massive maternal mRNA degradation and ZGA occur, suggesting that Nanos-mediated downregulation of maternal mRNAs is likely a part of the mechanism promoting MZT. In *C. elegans* PGCs, Nanos has been reported to facilitate MZT by downregulating maternal mRNAs during a transcriptionally repressed state^2^. Thus, investigating the function of Nanos will provide important perspectives for understanding the MZT in PGCs across various animals.

Nanos-mediated mRNA downregulation under transcriptional repression occurs even in animals in which MZT does not occur in PGCs. In small micromeres of sea urchins (germline progenitors of *Strongylocentrotus purpuratus*), MZT does not occur; however, these cells maintain transcriptional repression during the blastula stage, and Nanos depletes certain mRNAs^23–25^. These findings suggest that, in animals such as *Drosophila, C. elegans*, and sea urchins, where the germline undergoes early development with transcriptional repression, Nanos-mediated downregulation of mRNAs is a common feature. Therefore, this process is likely to play an important role during the early stages of development.

## 3 EXPERIMENTAL PROCEDURES

### 3.1 Fly strains

Flies were maintained on standard *Drosophila* medium at 25°C. The following fly stocks were used: *y w, nanos*^*BN*9^, *EGFP-vasa*^26^. In the RNA-seq analysis, embryos produced from *nanos*^*BN*^ *EGFP-vasa*/*+* and *nanos*^*BN*^ *EGFP-vasa*/*nanos*^*BN*^ females mated with *y w* were referred to as normal and *nanos*^−^, respectively. In the RT-qPCR analysis, embryos produced from *EGFP-vasa /+*; *nanos*^*BN*^/*TM3* and *EGFP-vasa /+*; *nanos*^*BN*^*/ nanos*^*BN*^ females mated with *y w* were referred to as normal and *nanos*^−^, respectively.

### 3.2 RNA-seq analysis

One hundred PGCs were isolated from 2 to 3 h AEL embryos using flow cytometry as described in a previous study^27^. cDNA was synthesized using the SMART-Seq v4 Ultra Low Input RNA Kit for Sequencing (Clontech, 634890) as described previously^15,28^. Nextera XT library creation and RNA-seq were performed at the University of Minnesota Genomics Center using a HiSeq 2500 platform (Illumina), and approximately 20 million reads per sample (50-bp paired-end reads) were obtained. Three biological replicates were used for each genotype. The corresponding raw read count data were deposited in the DNA Data Bank of Japan (DDBJ) under accession number DRR720111–DRR720116.

Raw reads were processed using the Trimmomatic software (ver. 0.36)^29^ and then aligned with HISAT2 (ver. 2.2.1)^30^ to the BDGP *D. melanogaster* genome (dm6). The SAMtools software (ver. 1.9)^31^ and StringTie (ver. 2.1.7)^32^ were used to sort, merge, and count the reads. Count reads were processed using prepDE.py3 (https://ccb.jhu.edu/software/stringtie/dl/prepDE.py3) and differential expression analysis was performed using the edgeR package (ver. 4.0.16)^33,34^ of R (ver. 4.3.3). For quality control of our dataset, the transcripts showing count-per-million > 0.5 in at least three samples were filtered and then normalized by the trimmed-mean method before differential expression analysis. Differentially expressed transcripts were identified by comparing the normal and *nanos*^−^ dataset using likelihood ratio test.

GO functional enrichment analysis was performed using Metascape (ver. 3.5)^35^. The detected transcripts were used as the background, and the significantly increased and decreased transcripts in *nanos*^−^ were analyzed separately.

To identify differentially expressed transcripts, the thresholds for statistical significance between normal and *nanos*^−^ PGCs were defined as FDR < 0.05 and log2FC ≥ 1 or ≤ −1. Subsequently, a list of all 3′-UTR sequences (FlyBase; dmel-all-three_prime_UTR-r6.61.fasta) was obtained. The 3′-UTR list was merged with the dataset of differentially expressed transcripts, and then the proportion and density of Nanos-Pum motif were calculated.

### 3.3 RT-qPCR analysis

One hundred PGCs were isolated from 2 to 3 h AEL embryos using flow cytometry as described in a previous study^27^. The isolated PGCs were sorted into PCR tubes preloaded with 4 μL of a lysis solution comprising RNase-free water (Takara Bio), 0.3% NP-40 (Thermo Fisher Scientific), and RNasin Plus at 1 U/μL (Promega). The tubes were mixed using MixMate (Eppendorf) at 2000 rpm for 30 s, followed by brief centrifugation at 4°C. cDNA was synthesized from the lysate using the Superscript VILO Master Mix (Thermo Fisher Scientific). Quantification was performed on a Light Cycler 480 system (Roche) using a QuantiTect SYBR Green RT-PCR Kit (QIAGEN). The primer sets used for qPCR are listed in Table S5. Thermal cycling conditions were as follows: one cycle of 95°C for 15 m, then 45 cycles of 95°C for 15 s and 60°C for 1 m. Fluorescence in each well was monitored throughout the cycling period. Melting curve analysis was performed to evaluate off-target amplification. Data were analyzed using the Light Cycler software (Roche) and Microsoft Excel (Microsoft). Using the ΔΔC_T_ method^36^, values were normalized against those of *Actin 5C* (*Act5C*), and log2 expression ratios were calculated. Statistical significance was calculated by paired *t*-test.

## ACKNOWLEDGEMENT

This work was supported in part by Grants-in-Aid for Scientific Research from the Japan Society for the Promotion of Science, Japan (JSPS) [KAKENHI Grant Numbers: 24H02030 (SK) and 23K05778 (MA)] and Japan Science and Technology Agency, Japan (JST) [Support for Pioneering Research Initiated by the Next Generation (SPRING) Grant Numbers: JPMJSP2124 (YK)]. We thank Dr. Shunta Yorimoto for technical advice of RNA-seq analysis.

